# Deriving general conditions and mechanisms for division of labor using the cell-based simulator gro

**DOI:** 10.1101/363093

**Authors:** Paula Gregorio-Godoy, Guillermo Pérez del Pulgar, Marcos Rodríguez-Regueira, Alfonso Rodríguez-Patón

## Abstract

Division of Labor can occur as a consequence of a major evolutionary transition such as multicellularity but is also found in societies of similar individuals like microbes. It has been defined as a process that occurs when cooperating individuals specialize to carry out specific tasks in a distributed manner. This paper analyzes the conditions for division of labor to emerge as a beneficial evolutionary solution and proposes two novel mechanisms for this process to emerge as a consequence of cell communication in an isogenic group of cells. The study is conducted by means of the cell-based model *gro* that simulates the growth and interaction of cells in a two-dimensional bacterial colony. When the labor is social, like the production of a molecule that is publicly shared, simulation results indicate that division of labor provides higher fitness than individual labor if the benefits of specialization are accelerating. Two genetic networks that generate consensual and reversible specialization are presented and characterized. In the proposed mechanisms, cells self-organize through the exchange of certain molecules and coordinate behaviors at the local level without the requirements of any fitness benefits. In addition, the proposed regulatory mechanisms are able to create *de novo* patterns unprecedented to this date that can scale with size.

## MAIN TEXT

Multicellularity is considered one of the major transitions in evolution, a concept that encompasses the phenomena in which natural selection transforms autonomously replicative genes/cells/individuals into new higher-level structures^1,2^. In such structures, individuals are no longer independent of each other but rather form a group that coordinates and communicates with a resulting behavior that is determined by the interaction of its fundamental elements^2^. One of this major transitions is multicellularity^3^. It has been identified that multicellularity has evolved independently more than 20 times, at least once in animals, three times in fungi, six times in algae, and multiple times in bacteria^4^.

Multicellularity has involved different evolutionary pathways, leading to the belief that there is not a single explanation for its origins. However, some requirements have been identified across many species and are thought to be essential to meet the current definition of what constitutes a multicellular organism^5^. The evolutionary requirements for multicellularity include (i) the formation of clusters of cells via physical contact or adhesion to form a new evolutionary unit^6^, and (ii) communication among the individuals leading to coordinated activity^7^.

There are some essential disadvantages to multicellularity. For example, it is much costly to adhere and communicate with your neighbors than to live a solitary life. Employing strategies that assure communication and encourage cooperation to reduce possible conflicts within the group does not come for free^8^. Yet, these pitfalls can be greatly compensated by the benefits that multicellularity can provide. It has been proved to improve: colonization of new territories^4^, predation avoidance^9^, resource acquisition, metabolic efficiency, higher resistance to physical and chemical stresses^10^ and, finally and more importantly, it implies cell differentiation and division of labor (DoL)^1^.

DoL requires the coexistence of interacting specialized individuals that carry out complementary tasks^11^. Accordingly, specialization refers to a process where the individual changes from a state where in principle many functions could be performed, to a configuration where it mainly develops a specific task^12^. The specialization of some individuals of the population usually implies that they become dependent on the rest of the population. Hence, individual cells renounced their capacity to reproduce as independent units and come to reproduce as part of a larger whole.

Several types of microbes have evolved some sort of DoL among its colony members and could therefore be considered multicellular organisms^13,14^. Bacterial expressions of DoL take different forms which range from undifferentiated chains to morphologically differentiated structures^10^. Some well-known examples include myxobacteria and cyanobacteria. Filamentous cyanobacteria are able to carry out two incompatible tasks, nitrogen fixation and photosynthesis, in two spatially separated and specialized cells: the carbon-fixating vegetative cells and nitrogen-fixating heterocysts^15^. In contrast, myxobacteria present an aggregative and motility‑dependent multicellularity. During starvation, myxobacteria undergo a dramatic transition where individual cells migrate together to create fruiting bodies for the spreading of the spores. Around 50% of the colony cells die during this transition in a process called programmed cell death (PCD), in order to supply nutrients for the fruiting body cells^16,17^.

It is natural, therefore, to assume that the requirements for multicellularity could, in principle, be met in a bacterial colony. Bacteria are social beings that because of their high reproductive rate do not disperse after division but rather stay together and form communities^18^. Cluster formation occurs in many biofilm structures and bacterial communication mechanisms are used to collaborate and coordinate different activities within the colony^19^. Bacteria often exhibit coordinated behavior focused on the survival of the colony (as in a multicellular organism), where they rely on the costly production of certain molecules that provide shared benefits for the community^20,21^. Examples of these secreted molecules range from informational signals, like quorum sensing^22^, to biofilm polymers^23^, iron-scavenging siderophores^24^ or digestive enzymes^25^.

This paper explores, through the simulation of bacterial colonies in various conditions, the subsequent conditions for DoL to emerge as a beneficial evolutionary solution. According to West & Cooper^26^, these conditions include: (i) DoL should provide a benefit for all individuals^26^, and, (ii) individuals should exhibit phenotypic variation and cooperation among them^27^. Results will be presented in two sections, where each section is related to one of the conditions mentioned above.

No specific bacterial strain is modelled, as simulations recreate general rules that can be found in most microbial environments. Simulations were performed using the cell-based model *gro*^28,29^. Cell-based models can be conceived as mechanistic representations of evolutionary games^30^ where the replication, interactions and death of individual agents are explicitly simulated using a system updated by a series of discrete events^31^. These models derive the global dynamics of the population from local interactions rules described for every single individual^32^. The behavior of the population then emerges because of the interaction between these individuals and the described environment. This feature is of crucial importance because it avoids the definition of the relation between the individual scale and the population level, which is normally required to be explicitly described by the fitness function of mathematical models^33^. The *gro* cell-based model is a two-dimensional representation of a biofilm, and comprises two entities: bacterial cells and its environment. The two-dimensional environment represents an infinite petri-dish discretized in equally-sized square units. The grid serves as a scaffold for the spatial localization of each cell. It also handles the diffusion of signaling and nutrient molecules. Bacterial cells are represented as rod-shape rigid bodies. The main state variable, that characterizes each cell, is an array of the protein expression levels associated to the bacterial genetic material. Proteins are digital abstractions that can only be in two states: expressed (1) and non-expressed (0). The behavior (e.g. growth rate, emission rate, nutrient uptake rate…) of every cell is then determined by the cell state, which is in turn, defined by the protein array. Following the qualitative approach of this paper, only the behavior of cells is considered, and therefore, proteins are bypassed and used only to connect cell state to its behavior.

## RESULTS

### DoL provides higher fitness if the benefits of task specialization are accelerating

Bacterial populations engage in social interactions through the emission of different diffusible molecules^34^. In order for DoL to make sense, the bacterial collective has to be required to perform, at least, two different tasks. Note that it is important that these tasks have to be social and not individual, as their products have to be shared by all. Only in this context could DoL provide any benefit. Simulations study when it is more beneficial for a population to divide the labor of two tasks compared to a situation where all individuals perform both tasks simultaneously. Results measure the performance of a population where both tasks are associated to the emission of diffusible molecules that, when present together in the environment, enhance cell growth. Populations with different degrees of DoL are compared (Figure 1). The main hypothesis, according to West & Cooper^26^, that motivates the following simulations is that the fitness benefit of DoL has to be accelerating. An accelerating fitness is recreated when the task becomes more efficient as more effort is put into it, or equivalently, when there is a penalty for the execution of incompatible tasks in the same location^35^. Simulations reveal which reward function produces an increase in the global fitness of the colony, which is measured as the number of cells after certain simulation time.

**Figure 1.**
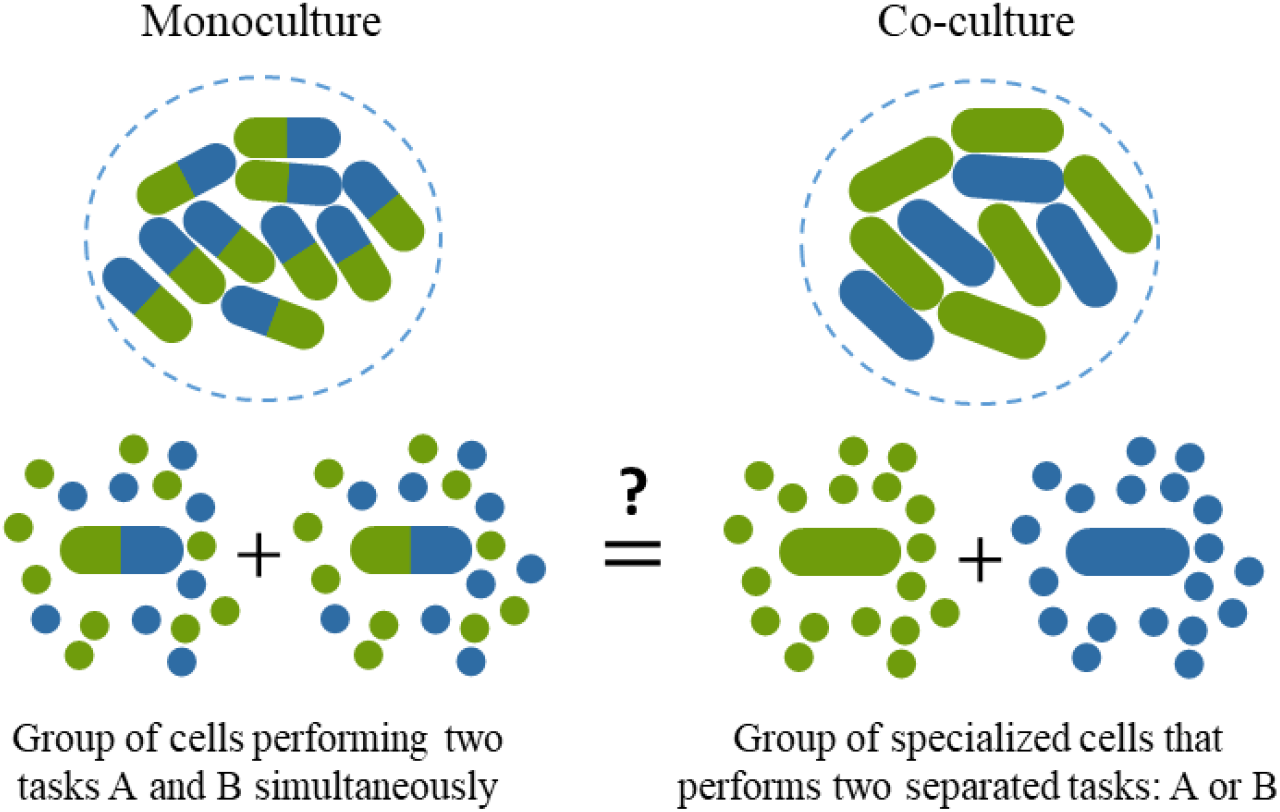
When is it more beneficial for a population to divide labor compared to all individuals performing two distinctive tasks? Because both tasks involve the emission of the product into the shared environment, it is possible that the whole group benefits from the labor without carrying the cost of production. The hypothesis is that the efficiency in the molecule emission should be greater in the co-culture case for DoL to be more beneficial than the monoculture.

Simulations consider a population of cells that performs two independent tasks. Both tasks involve the production and emission of a certain diffusible molecule, namely, molecule A (associated to task A) and molecule B (associated to task B). All cells are required both molecules to properly grow and divide. Each cell has a constant and finite set of resources to spend in the production of molecules. The imposition is that if any cell invests a proportion of resources (X) into task A, then, the remaining proportion (1-X) would be invested into task B.

The next assumption is that the benefits of DoL are going to depend on the degree of communication, which is given by the diffusion and degradation rates of the molecules. The diffusion and degradation rates of the shared molecules define a specific length of communication. This is, provided that a certain cell is emitting, for instance, one molecule A per simulation step into the shared environment, then, the communication length is given by the average distance where individuals can receive that molecule (Figure 2A).

**Figure 2.**
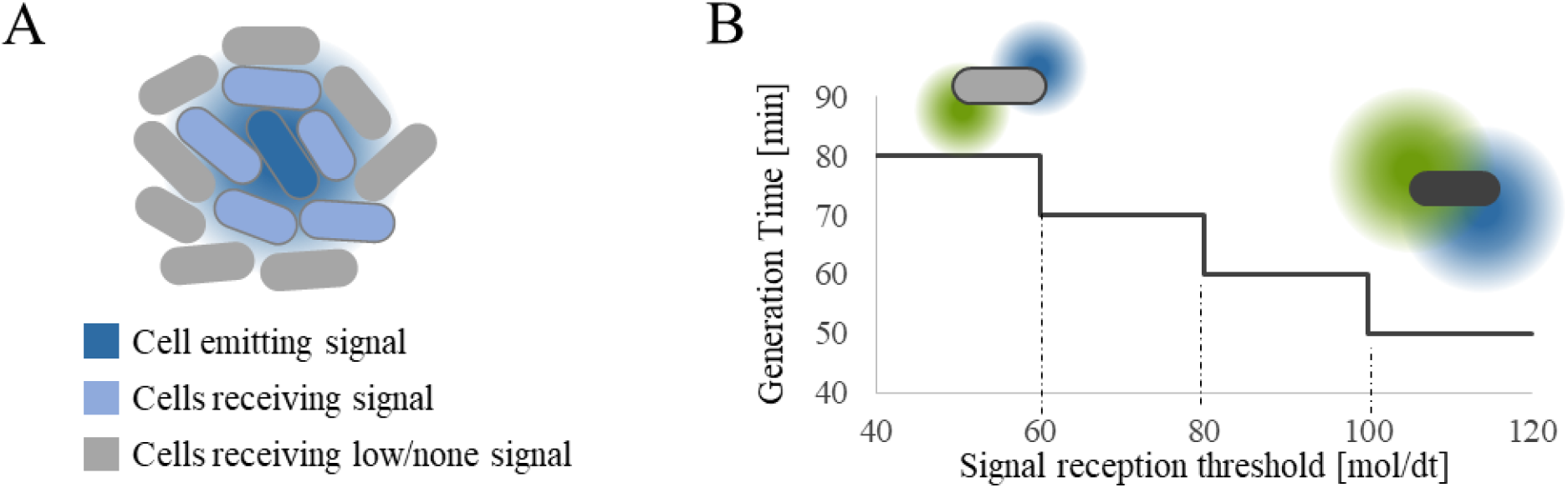
(A) Diffusion and degradation rates of the shared molecules establish the length of communication. With a diffusion of 0.2 (mol∙cell_size)/dt and a degradation rate of 0.3 mol/dt, a cell can communicate at least with its closest neighbors. The blue cell is emitting 1 molecule per time step and the reception threshold is set to 1 molecule per time step. Cells that sense above the reception threshold are depicted in light blue. **(B) Molecule reception dictates cell generation time.** The reception threshold of each cell modulates the cell individual fitness. This is, the division time of the cell depends on the amount of signal (of both A and B) that it reads from the environment. The more signal a cell can receive, the shorter the division time, and thus, the higher its reproductive fitness.

The individual fitness is dictated by the amount of signal that each cell can perceive (Figure 2B). The more molecules (of both A and B) a cell can sense, the shorter the generation time of such cell. The individual fitness of each cell, will translate into the global fitness of the colony as the total number of cells. This is the metric used for evaluating the benefits of each strategy. The only free parameter left is the emission rate of each cell. The model assumes that, for the same resources, the task (*i.e.* producing molecule) can be more or less efficient. This is modelled with different reward/cost functions that determine the emission rate of each cell, depending on its specialization status.

Simulations compare the overall fitness of the population after 4 hours of simulated experimental time. Each simulation consists on a colony composed by two subpopulations randomly distributed in space. Each subpopulation has the same initial number of cells (1000 initial cells per population, 2000 in total). The key matter is that the level of specialization of each individual within the subpopulation varies, ranging from subpopulations of non-specialized cells to subpopulations that only undertake one of the tasks. Simulations were run for the six different specialization scenarios (Figure 3).

**Figure 3.**
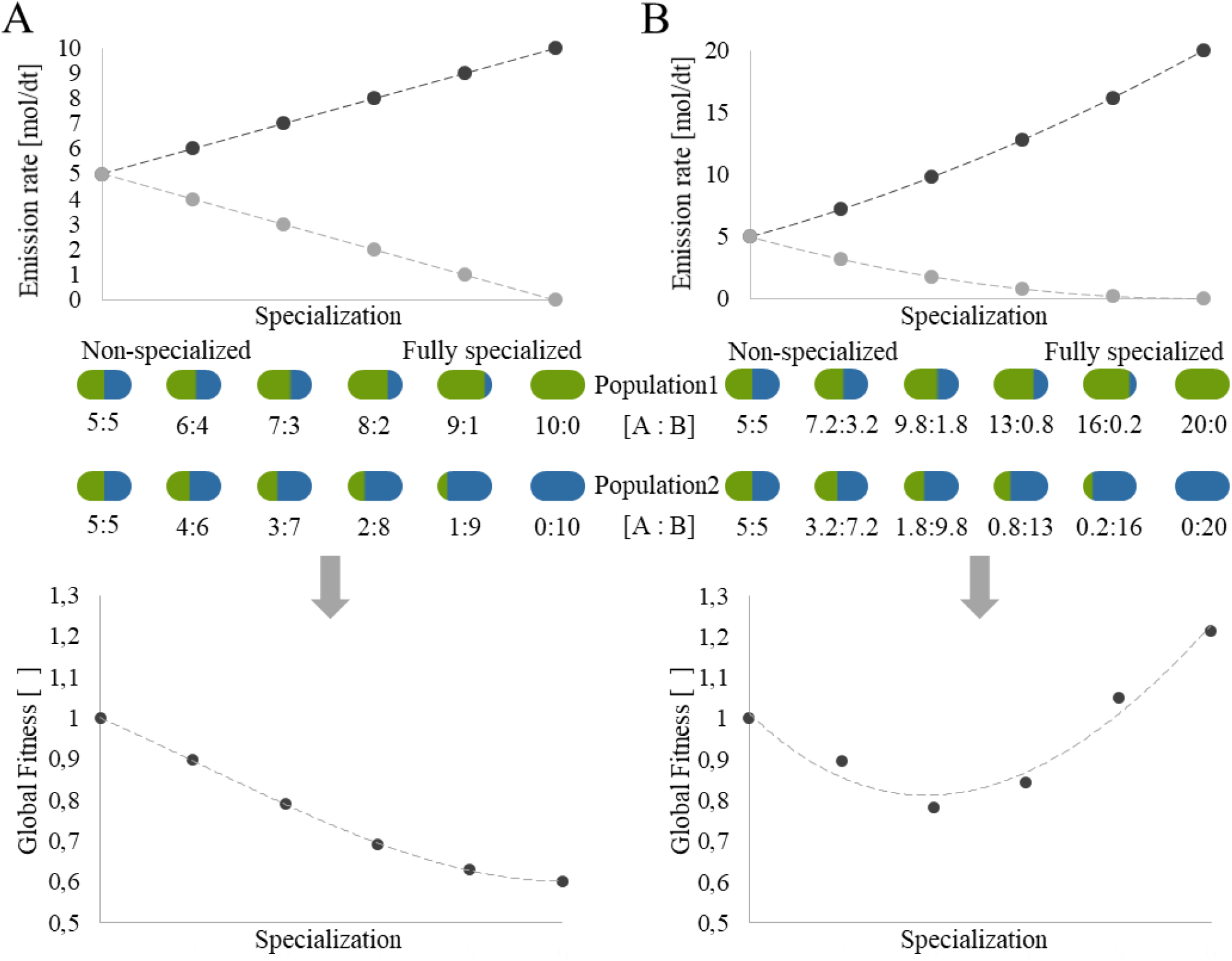
(A) Linear emission rates (top) of molecules A and B for six different degrees of specialization in subpopulation 1 and 2, and their corresponding global fitness (bottom). Statistical significance was achieved for N= 10 runs per scenario with negligible variance. The large initial number of cells removes the global stochasticity that could arise, proving that the global fitness as defined (number of cells after 4 hours of experiment time) is a reliable and robust readout. The final number of cells in the non-specialized experiment is used as a reference value for normalization. The average global fitness declines as cell specialize. Notice that total specialization returns a global fitness that is almost half of the non-specialized case. This proves that a constant cost function does not promote DoL over individual production. **(B) Exponential emission rates (top) of molecules A and B for six different degrees of specialization in subpopulation 1 and 2, and their corresponding global fitness (bottom).** The global fitness initially decreases for cases with partial specialization where the reward in signal emission does not compensate still the loss by diffusion. As specialization is more extreme, the global fitness increases. Not complete specialization (16.2:0.2) already returns a higher fitness than the non-specialized case (global fitness of 1.1). Finally, fully specialization achieves an average fitness 1.3 higher than the non-specialized case proving that accelerating benefits, in the form of more efficient molecule production, are required for DoL to be a successful strategy.

In Figure 3A, the reward/cost function is translated into a linear emission rate (top figure). The first scenario considers two equal subpopulations that do not divide labor, where their cells simultaneously produce molecules A and B. Each cell has an emission rate of 5 molecules of each type per time step. Therefore, at each time step, 2000 cells are emitting 5 molecules of A and 5 molecules of B. The next experiment considers a case where cells are slightly specialized. In this case, population 1 consists of 1000 cells that have an emission rate of 6 molecules of A per time step while the emission rate of molecule B is set to 4. On the other hand, the remaining subpopulation, population 2, consists of 1000 cells that emits 4 molecules of A and 6 molecules of B per time step. Notice that population 2 is always complementary to the distribution of tasks of population 1, in all simulated scenarios. Overall, in all six scenarios, the global emission of each molecule is set to be 10 molecules per time step. Succeeding simulations consider individuals that are increasingly more specialized. The final experiment simulates the extreme case where subpopulation 1 consists only on cells that emit molecule A while cells of subpopulation 2 emit molecule B. Again, the global emission rate is kept constant and set to 10 molecules per time step. So, if the reception thresholds determine the individual fitness through the reproductive rate (Figure 2B), the emission rate defines the reward/cost of producing molecules A and B. Since the costs are constant and equal for all cells independently of their degree of specialization, the number of emitted molecules only depends on the amount of resources dedicated to each task, being this relation linear.

The obtained global fitness for this linear reward/cost function is shown in the bottom of Figure 3A. The final number of cells in the non-specialized experiment is used as a reference value for normalization. Notice that total specialization returns a global fitness that is almost half of the non-specialized case. This proves that a constant reward function does not promote DoL over individual production. The lack of benefit for specialization is a consequence of the molecule diffusion and degradation. Cells that produce both molecules are not required to wait for the already-diffused-and-degraded molecule that is coming from their neighbors. Since the intangible cost of diffusion associated to the process of specialization is not compensated in any way, there is no advantage for DoL in this context.

According to the initial hypothesis, an accelerating reward/cost function is also simulated. In the top of Figure 3B, the emission rate of each molecule is represented. The reward/cost function is now a function of cell specialization. Here, the cost per emitted molecule is reduced in proportion to cell specialization as the emission function represents the idea that the task becomes more efficient as more resources are put into it. The specialization in the production of a molecule is rewarded by increasing the rate of the emitted molecules for the same invested resource. That results in the emission of 20 molecules per simulation step for the fully specialized cell (Figure 3B, top).

The obtained global fitness for the exponential reward/cost function is depicted in the bottom of Figure 3B. The shape of the global fitness is a consequence of the trade-off that exists between the molecule loss due to diffusion and the higher efficiency that DoL provides. Results show how the global fitness initially decreases for cases with partial specialization where molecule losses by diffusion and degradation are still not compensated by the increase in production efficiency. These simulations provide a computational proof of the efficiency requirements for DoL to be a successful strategy.

Simulations are now extended to different emission functions (reward/cost functions) other than the linear and quadratic. A cubic emission function (∼linear_emission^3^) and the square root function (∼linear_emission^1/2^) are simulated for different degrees of specialization (central graph, Figure 4). The cubic emission function corresponds to a scenario where specialization is highly rewarded, while the square root function mimics a scenario where specialization is punished.

**Figure 4.**
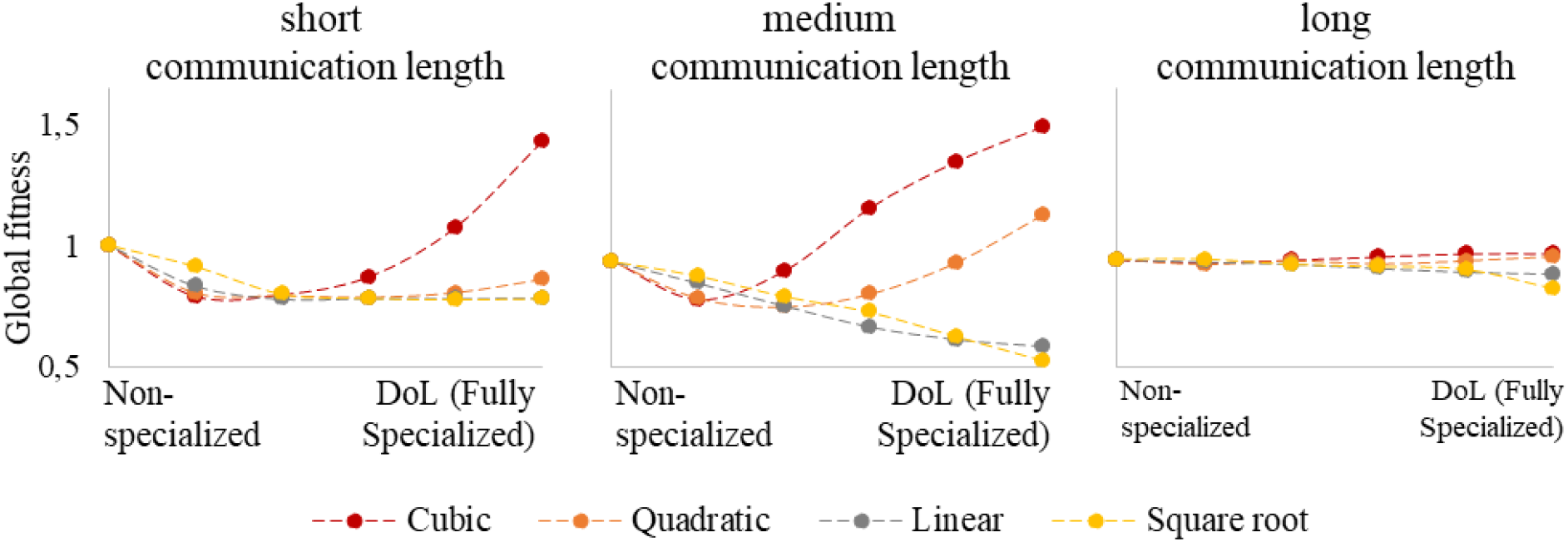
Global fitness with different degrees of specialization for different emission functions and different communication lengths. Results corresponding to the same emission function were connected with dotted lines for an easier visualization. The small variation between the different runs (N=10) leads to the substitution of error bars for big dots for the same reason. **Short communication length** (left graph) is recreated with a diffusion of 0.1 (mol∙cell_size)/dt and a degradation rate of 0.5 mol/dt. Molecules can only be sensed by the cell that is emitting them and occasionally by some close neighbors. In this scenario, only the cubic function is able to overcome the shortage in signal and rescue DoL populations that are fully specialized. **Medium communication length** (central graph) is recreated with the same parameters as Figure 2, diffusion of 0.2 (mol∙cell_size)/dt and a degradation rate of 0.3 mol/dt. Only accelerated benefits (quadratic and cubic) are able to increase the global fitness of a population that has divided labor. **Long communication length** (right graph) is recreated with a diffusion of 0.5 (mol∙cell_size)/dt and a degradation rate of 0.1 mol/dt. The range of communication is widened and the system becomes so saturated that accelerated emission functions do no longer provide a competitive advantage nor a disadvantage.

The central graph of Figure 4 depicts the global fitness for four reward/cost functions. Only accelerated functions (quadratic and cubic) are able to increase the global fitness of a population that has divided labor. Notice that for slight specialization the constant and the square root function return higher fitness than the quadratic function. It is interesting that the square root function, which implies decelerated benefits or in other words less emission in the specialized case than in the non-specialized, returns the same fitness tendency than the linear function.

It is important to notice that the benefits of DoL can emerge only if individuals are able to communicate with to each other. The effect that the degree of communication has in promoting DoL is considered as well in Figure 4. The degree of communication, which is represented as the length of communication, is changed in the following simulations to include shorter and longer communication lengths. Results in the short communication range are shown in the left graph of Figure 4. Short communication length (diffusion=0.1 mol/dt; degradation=0.5 mol/dt) simulates a signal that can only be sensed by the cell that is emitting it and occasionally by some close neighbors. Because the half-life of that molecule is so small, the accelerated benefits of the quadratic function cannot compensate the loss of signal, and DoL results as a poorer strategy than non-specialization. The general fitness of specialized populations is lowered as they rely on a more long-live communication system. It is only the cubic function that is able to rescue specialized populations by producing much more signal. On the other hand, for long communication lengths (Figure 4, right graph) when the range of communication is widened (diffusion=0.5 mol/dt; degradation=0.1 mol/dt), the system becomes so saturated that accelerated emission functions do no longer provide a competitive advantage. The half-life of the molecules is so high that there is no loss in information when sharing, and the specialized and non-specialized populations become almost undifferentiable for all emission functions.

### Cell coordination can generate DoL through the emission and sensing of diffusible molecules in homogeneous populations

Even in homogeneous environments and isogenic populations, bacteria can exhibit cell-to-cell phenotypic variability^36^. This is referred to as individuality^37^. Phenotypic heterogeneity allows colonies to adapt and survive to sudden changes in the environment and is central in the emergence of DoL. The fundamental idea is that phenotypic heterogeneity can provide such benefits because, heterogeneous colonies have more diverse strategies to face the environment, and thus, are better adapted for evolution^38^. The most relevant internal molecular mechanisms that give rise to phenotypic variation are genetic differences, mutations or noise in gene expression^39^. For its emergence, DoL is said to require individuals to exhibit phenotypic variation, where not all individuals follow the same strategy. Here, two novel genetic circuits are explored as difference generators in an initially homogeneous population. Simulations study the ability of an initially undifferentiated population of cells to reach a consensus for labor distribution. The proposed circuits focused on the regulatory aspects of DoL and can be implemented in a variety of contexts. Both circuits are based on intercellular signal regulation, the first uses only one signal while the second regulates the specialization of the two tasks using two different signals. Exploration of the circuit properties is conducted mainly without any cost/fitness functions in stationary non-growing populations. Finally, their attributes as morphogenetic/patterning networks are also studied through their ability to scale, and their dependency with size and shape.

Following the context of previous simulations, a bacterial population that is required to perform to different tasks is simulated. All cells are equal and behave according to the rules that the internal circuit dictates. Here, cells do not perform the two tasks simultaneously but rather transit between two different states, A and B, each one in charge of carrying out a task. Unlike previous simulations, the behavior of each cell (its degree of specialization) is not imposed, but rather changes according to the cell internal logic and its environment.

The first mechanism is shown in Figure 5A. Tasks could be associated to one of both possible states, A or B. In this case, the task associated to state B is the molecule emission while no task is associated to cells in state A. The molecule secreted by cells in the B state acts as a regulator to switch between the A and B states. Notice that here the emitted molecule has a regulating function for the emergence and maintenance of DoL. Cells can be either in the A state or in the B state, but not both. Each state has an associated sensitivity threshold. If a cell in the B state senses that the concentration of signal is higher than its threshold, then the cell transits to the A state. On the other hand, if a cell in the A state senses that the signal concentration is below its threshold, the cell would turn into the B state. Simulated cells begin in the A state and the lack of signal automatically produces all cells to turn into the B state. Then, the accumulation of signal converts some cells back to the A state until eventually a balance between the proportion of A and B cells is reached (Fig 5B).

**Figure 5.**
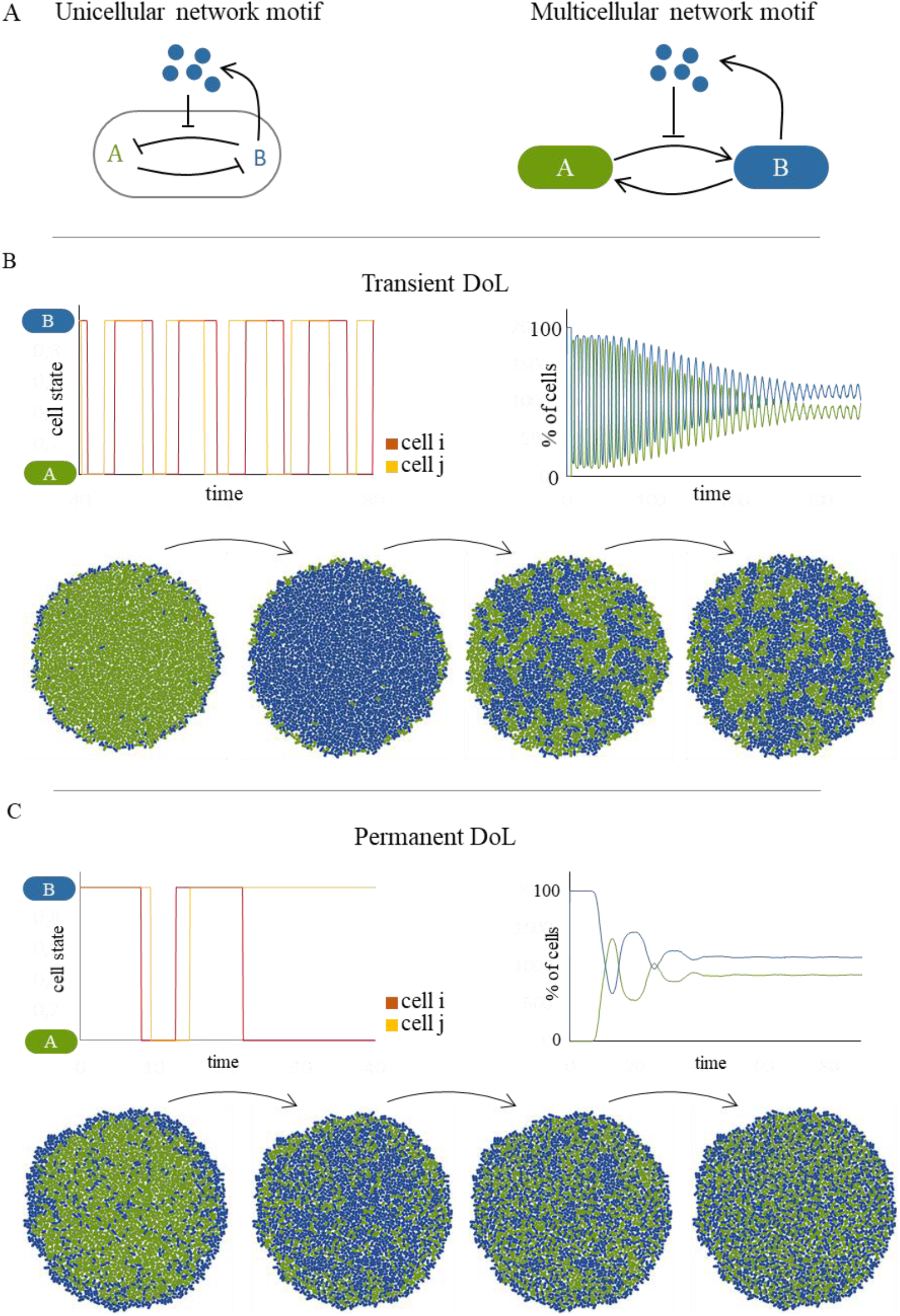
(A) Representation of the one-signal coordinated signaling circuit and its multicellular implications. The unicellular motif shows the inner states A and B and how they relate. Following the described rules, state A and B are mutually exclusive, hence the double repression. This represents the premise that cells can only be in one of the two states. The presence of the molecule in the environment weakens the repression of B over A, inducing the A state. The absence of signal reinforces the repression of B over A, and thus, cells turn into the B state. The multicellular motif shows the influence between neighboring cells through the emission of the regulatory molecule in the shared environment. Cells in B influence their neighbors (and themselves) by promoting the A state through the emission of the regulatory molecule. On the other hand, cells in the A state will passively promote the B state in their vicinity by not emitting the regulatory signal. **(B) Transient DoL**. The internal transitions of two random cells (i, j) show how cells oscillate between state A and B with different periods. The graph on the right depicts the number of cells in each state over time where the blue line represent cells in the B state and the green line represent cells in the A state. Temporal shots of the bacterial colony depict cells oscillating between the A state (green) and the B state (blue). Cells oscillate between the two states, at first synchronously and then increasingly asynchronously. The initial synchrony is visually observed in the shots, as all cells are in one of the two states. As the signal diffuses over the colony, desynchronization starts to happen first at the edge, and then later spreads all over the colony. Eventually, the number of cells in each state converges into a certain proportion. Specialization is achieved in a transient manner with 60% of the population performing task B and the remaining 40% performing task A. Despite this convergence, the specific individuals that carry out each task changes with time. **(C) Permanent DoL.** The internal states of two random cells shows how in this regime cells rapidly fixate in one of the two states permanently. All cells end up in a given state and remain there indefinitely. However, each cell has a specific differentiation time, as cell j stabilizes in state B after one oscillation while cell i requires two periods to stabilize in state A.

The behavior of the whole system depends on the model parameters, namely, the diffusion and degradation rates of the coordination signal, the signal emission rate and the sensitivity threshold of both states. An exhaustive exploration of the parameter space study has been conducted and can be found in the SI (Figure S1). This circuit is able to generate five different behaviors, depending on the model parameters, that include transient DoL (Figure 5B), permanent DoL (Figure 5C), global oscillations between state A and B, and also, cells can get stuck in either state permanently. DoL is said to be transient if cells switch between states perpetually, but the number of cells in each state converges into a certain proportion. In Figure 5B, DoL is achieved in a transient manner with 60% of the population performing task B and the remaining 40% performing task A, on average. On the other hand, DoL is said to be permanent if cells end up in only one of the two states resulting in a static solution. Figure 5C depicts how permanent DoL is achieved. After a few initial oscillations the system stabilizes, and permanent DOL emerges, with cells stuck in the state dictated by the states of their neighbors. This is due to signal coordination among neighboring cells. Because of the asynchrony among neighboring cells, there is a chance that a cell meets its requirements for the regulating molecule by producing it or by being near a producer cell, losing the need to change its state in the future. Cells do not engage in this behavior at the same time, but rather, the coordination is performed in small groups that communicate with each other in a domino-like fashion.

Surprising spatial patterns appear for a certain range of parameters (Figure 6). In these simulations, the proposed motif was able to achieve multicellular consensus and permanent DoL. As in the previous case, cells fixate on a certain state that accommodates the local need of the regulatory signal. However, unlike simulations from Figure 5C which returned a scattered distribution of cells of both states, here, cells organize in space creating ordered patterns.

**Figure 6.**
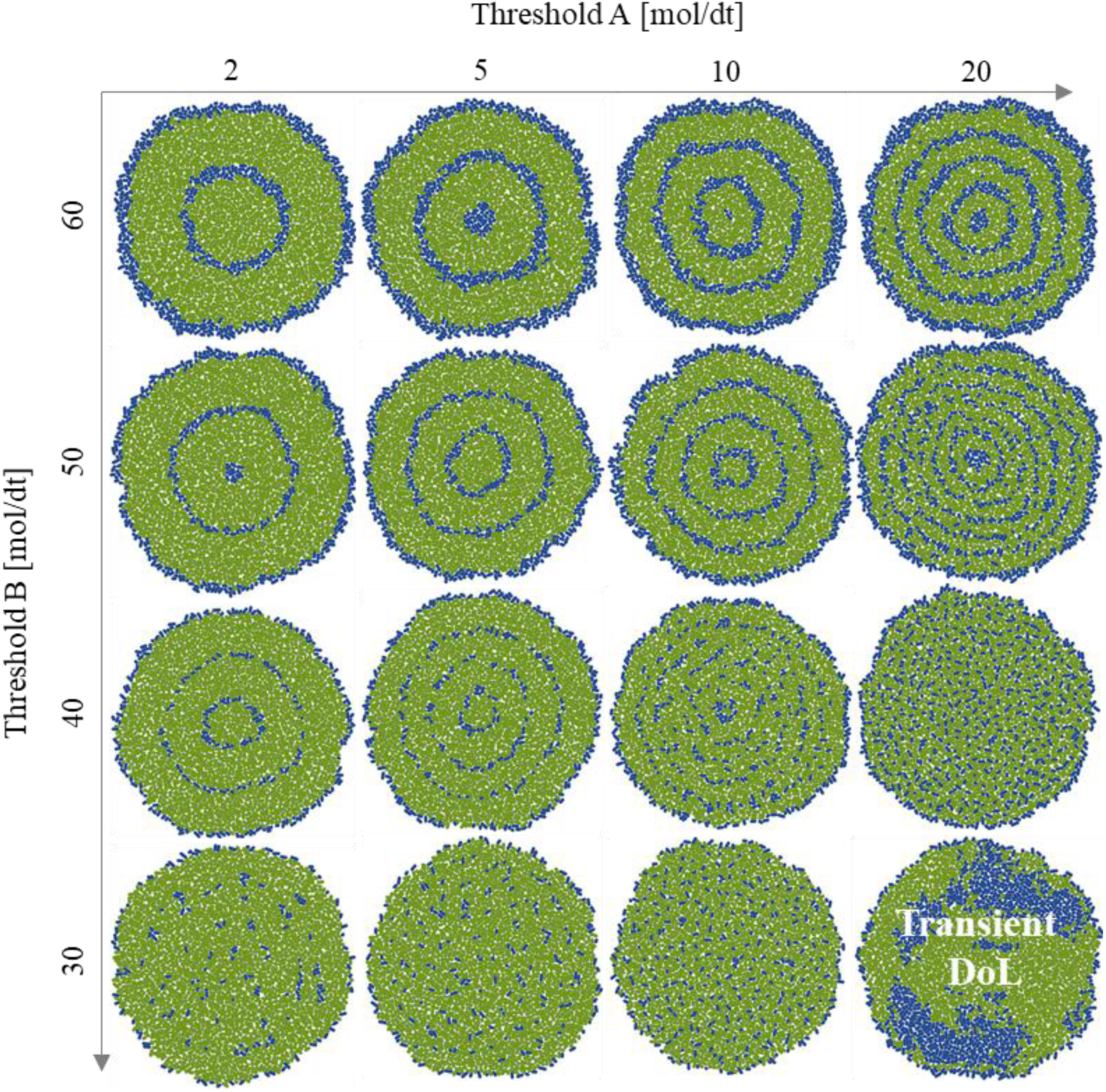
Spatial patterns emerge as a consequence of cell specialization. The threshold for B controls the width of the blue ring while threshold for A controls the width of the blue ring, and thus, the number of blue rings per colony. Cells fixate in the B state (blue cells) in the border of the colony because the signal concentration is lower in that region, activating the sensitivity threshold of the B state. The fixation of cells in the edge in the B state induce the A state in their inner neighbors. This fixation of the A state triggers, on a later stage, a second ring of cells in the B state. This cascades until the center of the colony, thus, creating the ring-like pattern.

The depicted range of values of threshold A (*i.e.* concentration of molecule below which a cell in state A turns to state B) and threshold B (i.e. concentration of molecule above which a cell in state B turns to state A) in Figure 6 corresponds to the values that exhibited permanent and ordered DoL. The only exception is the bottom right image, which was not able to stabilize neither in time nor in space. Notice that the green region is where the signal concentration is higher than the A threshold, and the blue region is where the signal concentration is lower than the B threshold. In a way, the A threshold acts as signal sink, and as it threshold increases, the regions where the signal concentration is not high enough are supplied by the emergence of B cells. The higher the threshold, therefore, the higher the requirement of B cells.

Now, the one-signal motif is extended by adding a second regulatory molecule. Inspired by the same logic, the purpose of this design is to achieve more robust and flexible behaviors. The internal and multicellular motif of the proposed circuit are shown in Figure 7A. The circuit imposes a series of rules that every individual in the colony must follow. The general functioning rules of this extended motif are:

- Cells have two possible states, A or B. Both states are mutually exclusive.
- Cells in A emit the diffusive molecule A whereas cells in B emit molecule B.
- Cells *work* to have both molecules A and B in their surroundings. Therefore, when individual cells sense that the concentration of the molecule they are not producing is below its sensitivity threshold and the one they produce is abundant, they change to the opposite state (e.g. if a cell in A senses that the concentration of molecule B is below its threshold while the molecule A is above its threshold, the cell would transit to state B).
- When the concentration of both A and B molecules is above the cell sensitivity threshold, cells will remain in its original state.
- When the concentration of both A and B molecules is below the cell sensitivity threshold, cells will remain in its original state.
- By convention, the system is initialized in the A state.

**Figure 7.**
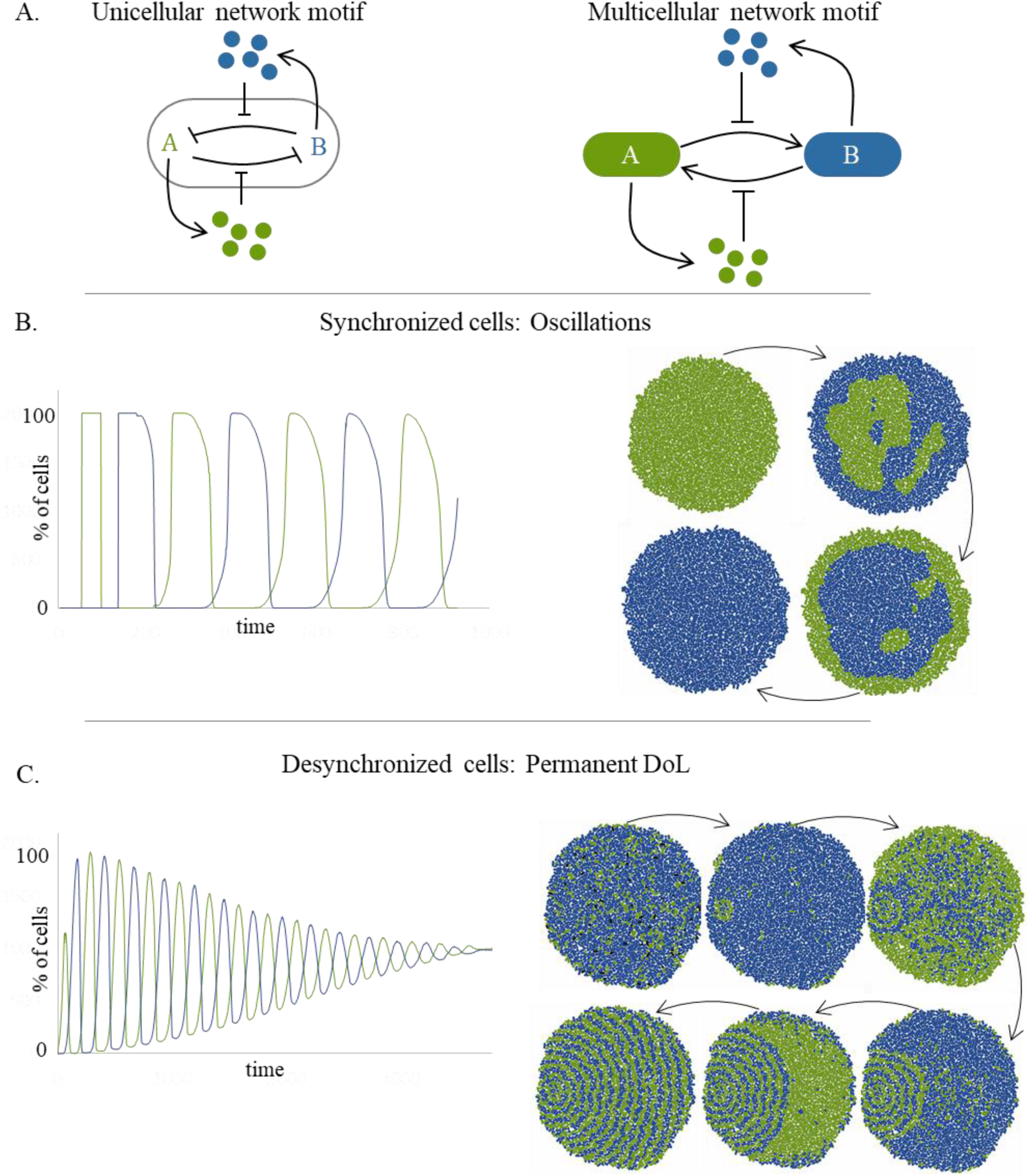
(A) Representation of the two-signals coordinated signaling circuit and its multicellular implications. The represented motif follows similar rules to the one presented in Figure 5 (cells in a certain state induce the opposite state in its neighbors), however here, both states have the associated task of emitting two different molecules **(B) Synchronized cells oscillate between state A and B.** The graph shows the number of cells in each state over time. The resulting global dynamics are oscillations of cells from state A (green) to B (blue). Temporal shots of a colony with the two-signal circuit depict how cells oscillate between the state A and B. Cells start by simultaneously entering the state A where they express the molecule A (represented in green in the figure). When enough molecule A has built up and the sensitivity threshold has been reached, cells commute to state B where they express molecule B (represented in blue). Because of the diffusive nature of the molecule, these transitions start in the edge of the colony and spreads towards the center. **(C) Desynchronized cells reach permanent DoL.** Cells engage in molecule sharing and divide labor in a permanent way. Since cells whose state has been locked influence other cells, the process spreads in the colony creating a very characteristic spatial pattern. The fraction of cells that engage in the process grows linear with time. Initially, cells start to oscillate between the state A and B to meet their molecule requirements. Randomly, a peak of one of the molecules somewhere in space induces that a cell fixes its state. In turn, this cell will induce the fixation of the opposite state in its neighbors, giving birth to a nucleation point. When the nucleation of the pattern starts, it starts to recruit cells that equally split the task of producing both molecules. Cells that are still not recruited continue to oscillate until they are influenced by their close neighbors. The oscillations in the number of cells start to lose amplitude as the pattern stablishes, and finally, it reaches a stable equilibrium where half the population is performing task A and the other half is performing task B, all of them in a precise spatial location.

Simulations explore the properties of the proposed motif in a static, non-growing population of cells. Model parameters in this circuit include the diffusion and degradation rates of molecules A and B, the cells emission rate of molecules A and B, and their respective sensitivity thresholds. Additionally, the production time of molecules is included as a parameter in this model. A full characterization can be found again in the SI. Simulations shown in Figure 7 consider the symmetric case where molecules A and B share the same attributes. Also, the sensitivity threshold and the emission rate of cells will also be the same for both A and B molecules.

Figure 7B depicts the colony behavior with this circuit when the production time of both molecules is noiseless. In this case, cells oscillate between the state A and B. Cells start by simultaneously entering the state A where they express the molecule A and when enough molecule A has built up and the sensitivity threshold has been reached, cells commute to state B where they express molecule B. Interestingly, if the production of molecules is not synchronized, a different phenomenon is observed. In Figure 7C, all parameters remain the same as in Figure 7B except for the production times of molecules A and B that now present a small variability (20 ± 5 minutes). The result is that cells do not longer oscillate but, after some time, get locked in a given state due to signal coordination among neighboring cells. Because of the time variability in molecule production, there is a chance that a cell meets its requirements for molecule A and B by producing, for example, molecule A and getting molecule B from its surrounding neighbors. This creates what is referred to as nucleation point. The generation of nucleation points was not possible in the previous simulation (Figure 7B) because cells were highly synchronized, and thus, all neighbors were roughly performing the same tasks at the same time.

The logic rules of the presented motif can be better exemplified with the following analogy. Imagine that the community of cells is a small village. Villagers need both water (molecule A) and bread (molecule B) to live, and they organize so that there is always enough water and bread for everybody. At first, everybody goes to the well to get water, and then, some will start baking bread, since there is none yet. The easiest way for the villagers is to share their product with their immediate neighbors. So, if a villager is currently baking bread he will need to get water from his neighbor. Yet, his neighbor has just starting baking bread because his neighbor across the street is collecting water. Then, the first villager will switch his task to water collection and stop making bread, as there is an excess of bread in his surroundings. Sometimes, with time, all villagers will get organized without an explicit general command from the mayor (i.e. transient and permanent DoL).

The first requirement for this motif to provide DoL is that neighboring cells need some kind of variability in their states. If the system is purely symmetrical and the production time of molecules is synchronized, cells will constantly change their state almost in unison. It would be like having only one cell permanently switching from task A to task B. This was not a requirement in the one-signal circuit because it was already asymmetrical by definition. Further characterization of the systems solutions is shown in Figure 8. Three different behaviors result from this exploration, where permanent DoL is achieve for almost all the parameter space. This proves that this motif is more robust than the one-signal design.

**Figure 8.**
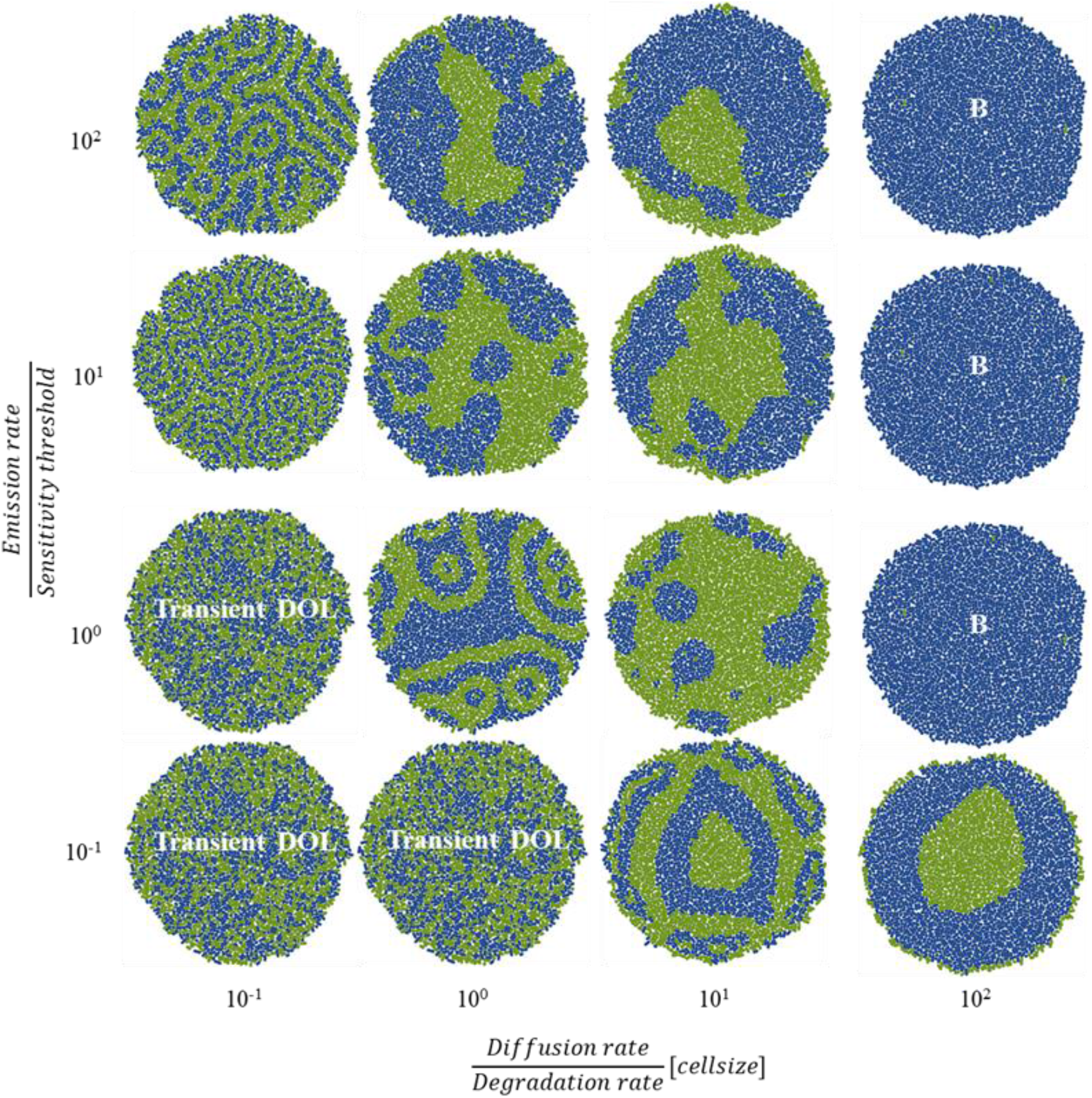
Resulting spatial patterns for different combinations of Emission rate/Sensitivity threshold and Diffusion/Degradation rates. Three different dynamics are obtained. DoL is achieved for all combinations except when the diffusion rate (1 (mol·cellsize)/dt) of both molecules is much higher than the degradation rate (0.01 mol/dt). In these cases (right column), the spatial scale of the signal is larger than the colony. Cells begin in state A (green) and the production of A saturates the environment. When cells turn to state B (blue) the same happens. Since cells are instructed to remain in the same state when both molecules are above their sensitivity threshold, they remain in the B state for as long the simulation lasted (300 minutes). In the remaining parameter space DoL is achieved. Transient DoL emerges when spatial scale of the signal is short and emission is lower than the sensitivity threshold. It is shown how when the emission is lower than the sensitivity threshold, cells form the salt & pepper pattern as it represents the smaller vicinity arrangement. In these cases, cells can only interact with their immediate neighbours because the regulating signal is both spatial and temporary short. Permanent DoL with ordered patterns appears when the spatial scale of the signal is short, but it is temporarily more stable. This happens when the emission rate is higher than the sensitivity threshold. The larger the difference between those values, the thicker the pattern. Finally, unordered patterns of cells undergoing permanent DoL are found for medium ranges of diffusion and degradation rates.

Independently of the pattern, permanent DoL in these simulations always provided an equal fraction of cells in state A and B (50-50). This is because the assigned parameters were considered equal for both states. Still, the fraction of cells in each state can be tuned to create any distribution (Figure S4 and Figure S5 of the SI show this effect).

The potential of the proposed genetic circuits may serve not only as difference generators for DoL, but it is also capable of creating spatial patterns interesting for morphogenesis. Morphogenesis is the biological process that causes an organism to develop its shape^40^. This process usually requires the pattering of cells via their spatial distribution during the embryonic development of an organism^41^. As it turns out, there are common mechanisms found in both DoL and morphogenesis, as seen, for example, in the patterning of nitrogen-fixing cells in cyanobacteria^15^. Both morphogenesis and DoL require some sort of specialization or differentiation among cells, but morphogenesis also involves this specialization to be ordered in space^42^. Some of the mechanisms involved in morphogenesis occur at the cellular level, with cell adhesion, proliferation and motility as mechanical forces that shape the final morphology of an organism^43^. Still, many morphogenetic processes occur at the genetic and molecular level, where the interaction of gen products and molecules (here named morphogens) are the cause of pattern formation^44,45^.

### Morphogenetic properties of the proposed mechanisms for the emergence of DoL

While many morphogenetic models based their spatial structure on pre-existing information, some interesting models of pattern formation are able to create patterns *de novo* in homogenous fields of cells with no existing cues^46^. An important mechanism for creating differences among initially homogeneous cells is lateral inhibition, a type of cell-cell interaction whereby a cell that adopts a particular state inhibits its immediate neighbors from adopting the same state^47,48^. This mutual inhibition mechanism between adjacent cells is usually mediated by Delta–Notch signaling system, as it happens in the Drosophila neurogenic ectoderm^49^. Another major example of self-regulated pattern formation is the reaction-diffusion model proposed by Turing as the basis for morphogenesis^50^. The model is based on two diffusible molecules that interact in a certain manner, leading to distinguishable ordered regions of each molecule.

These two important models of morphogenesis are mentioned here as they have served as inspiration for the proposed self-coordinating circuits. Notice that, in general, both the one-signal and the two-signal circuits are also self-regulating and do not need prior cues for the formation of spatial patterns, which is extraordinary. First of all, both proposed circuits employ some sort of lateral inhibition. In these circuits, where differentiation between two exclusive states is emergent, cells would not inhibit its state on neighboring cells but instead, activate the opposite mutually exclusive state. This interaction is not contact mediated as in lateral inhibition systems, but rather is mediated through the regulatory diffusive molecules. On the other hand, the basis of the Turing reaction-diffusive model is that two incoherent pathways are initiated at different speeds: a short-term direct or indirect self-induction is required, as well as, a long-term self-repression^44,51^. Both proposed circuits utilize this same logic. In the one-signal circuit (Figure 5A), as seen from the B state, the state self-induces by repressing its inhibitor state, A, within a cell (short-term self-induction). On the other hand, incoherently, B also self-represses through the emission of the diffusible molecule, which, in turn, weakens the B repression over A (long-term self-repression). In the two-signal circuit (Figure 7A), this logic applies both from the point of view of A and B states. This could be the reason for its robustness, since the Turing logical rules takes place symmetrically and simultaneously.

The plethora of spatial patterns are characterized based on two selected qualities: the number of nucleation points, and the *thickness* of the pattern. As previously commented, nucleation points are those cells that locally arrange and lock their state independently of the rest of the colony. They are responsible for the origination and propagation of the pattern and cannot be directed, as they are a consequence of a random emergent self-coordinating process. The *thickness* of the pattern refers to the average number of cells that lie between two equal-performing task regions. The magnitudes of the *thickness* and the number of nucleation points are used as reference to determine the dependency with size and shape of the representative patterns shown in Figure 9. The change in the pattering magnitudes with size as the group of cells grow is called scaling and is tested in Figure 10.

**Figure 9.**
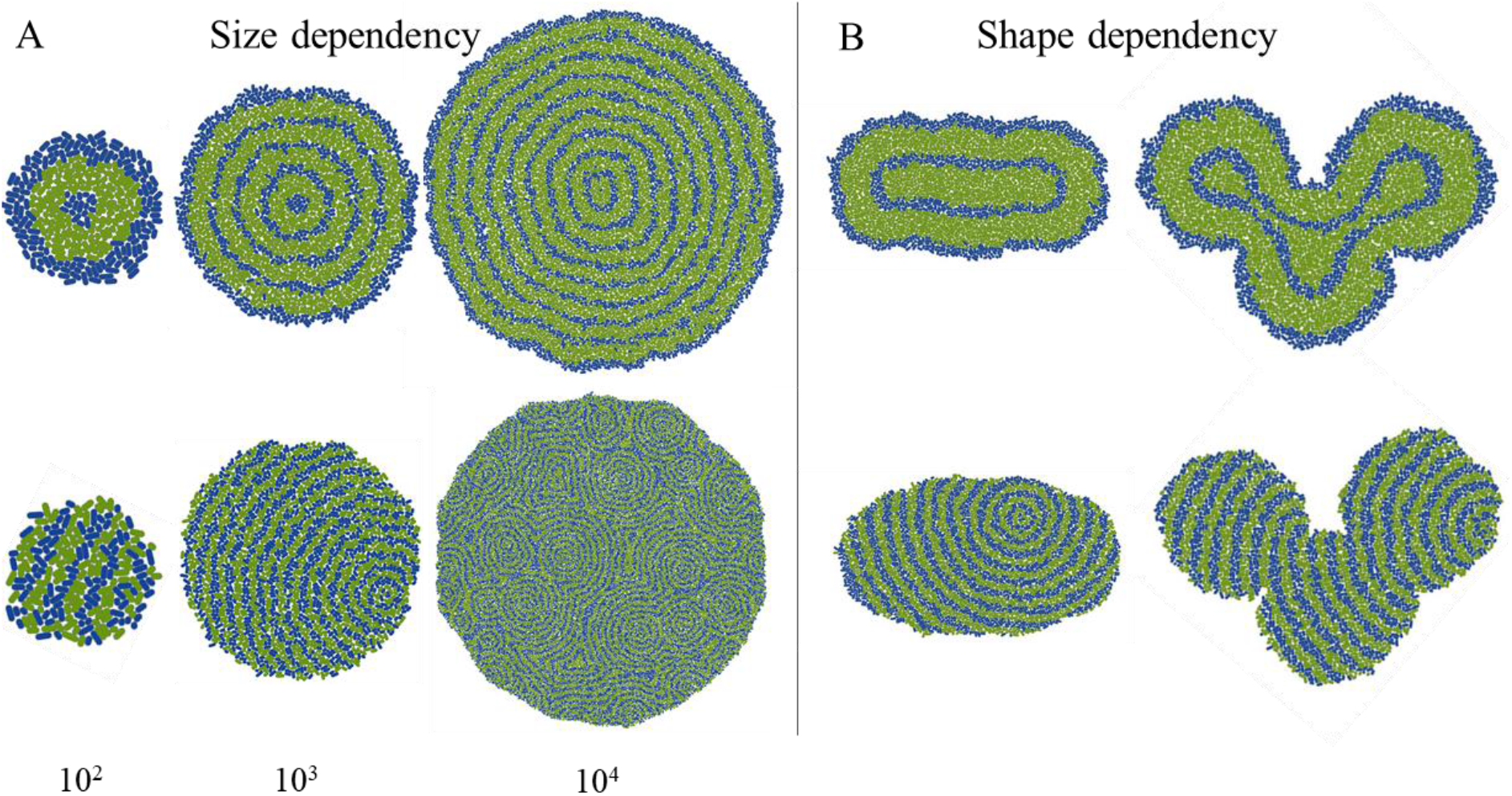
Pattern size (A) and shape (B) dependency. Each simulation was run for N=20 times, and the average value of the reference magnitudes was used as indicator of pattern variation between the different initial conditions. Nucleation points only apply in the two-signal patterns (bottom row). **(A) In the size dependency** study, simulations are initialized with three different number of cells, namely, 10^2^, 10^3^ and 10^4^ cells. The obtained *thickness* of the A region (green) is referred to as T_G_, while the *thickness* of the B region (blue) is referred to as T_B_. Results of the one-signal patterns (top row) for 10^2^ returned a T_G_=10.0±0.1 [cells], for 10^3^ a T_G_=10.0±0.1 [cells] and T_B_=2.0±0.1 [cells], and for 10^4^ a T_G_=10.0±0.1 [cells] and T_B_=2.0±0.1 [cells]. The blue *thickness* is also conserved for the 10^3^ and 10^4^ simulations although there is not enough space for a ring in the 10^2^ colony to compare with. Results of the two-signal patterns (bottom row) returned a T_G_=2.0±0.1 [cells] and T_B_=2.0±0.1 [cells] for all sizes. Nucleation points, *N*, varied according to the number of cells: *N*=1.05±0.01 (10^2^), *N*=1.7±0.4 (10^3^), *N*=15±5 (10^4^). **(B) In the shape dependency** study, simulations are initialized with two different bacterial dispositions. Results for the one-signal patterns (top row) for the elongated shape returned a T_G_=10.0±0.1 [cells] and T_B_=2.0±0.1 [cells]. The *thickness* of the heart-shaped colonies returned T_G_=7.5±0.1 [cells] and T_B_=2.0±0.1 [cells]. Results for the two-signal patterns (bottom row) for the elongated shape returned a T_G_=2.0±0.1 [cells] and T_B_=2.0±0.1 [cells]. The *thickness* of the heart-shaped colonies returned T_G_=2.0±0.1 [cells] and T_B_=2.0±0.1 [cells]. Both shapes showed the same average number of nucleation points, *N*=1.7±0.4.

**Figure 10.**
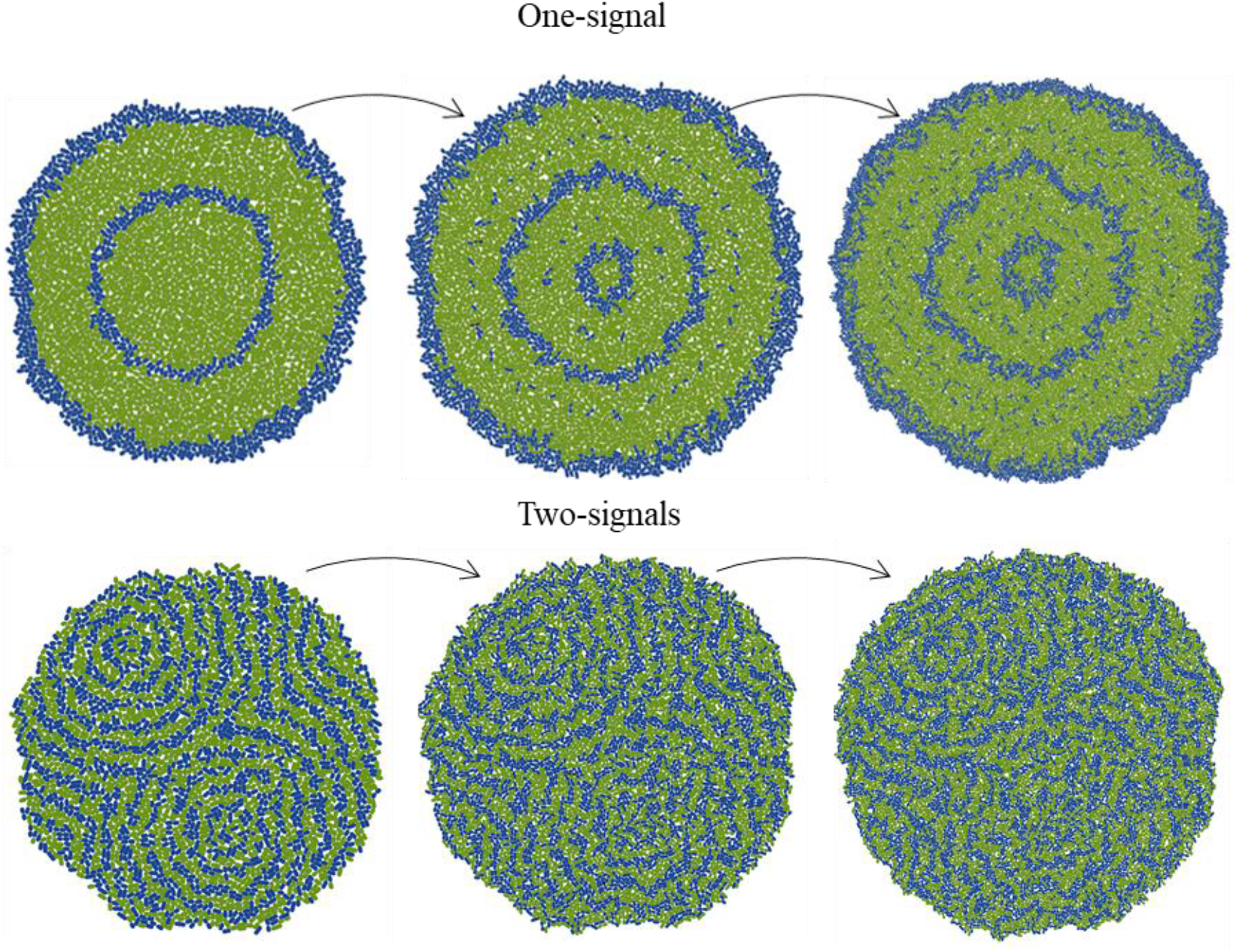
Pattern scaling of the one and two-signal circuits. Since cells growth rate is slower or similar to the temporal scale of the signal, the pattern scales accordingly although it blurs with size. The *thickness* is conserved throughout the entire simulation, with T_G_=10.0±0.1 [cells] and T_B_=2.0±0.1 [cells] for the one-signal and T_G_=2.0±0.1 [cells] and T_B_=2.0±0.1 for the two-signal patterns.

Size dependency refers to the dependence of the spatial pattern with the total body size, which in this case is the number of cells. The one-signal patterns (Figure 9A, top row) present equal *thickness* of the green rings. The blue *thickness* is also conserved for the 10^3^ and 10^4^ simulations although there is not enough space for a ring in the 10^2^ colony to compare with. In general, the ring-like motif resizes as expected for the different number of cells, revealing that the pattern is size invariant. The two-signal patterns (Figure 9A, bottom row) shows that the relative *thickness* is conserved across the various initial number of cells. The pattern general design changes due to the randomness (in number and location) of the nucleation points. The location of the nucleation points is a statistical phenomenon, while their number ultimately depends on the quantity of cells in a non-linear manner (Figure 9A).

Similar to the size dependency, the shape dependency refers to the correlation between the general shape of the colony and the imprinted pattern. Again, the reference magnitudes included in the caption of Figure 9 are referred to the average *thickness,* and number of nucleation points for each of the shapes. In Figure 9B, it is shown how even if variation between runs is low, the one-signal patterns (top row) are strongly influenced by the general contour of the colony. While the *thickness* of the blue region is conserved, the green inner region adapts to the spatial solution and varies strongly between shapes. On the other hand, the two-signal patterns (bottom row) show that despite the low repeatability due to the randomness of the nucleation points, its average value was not affected by the colony general shape. The *thickness* is also conserved, proving that these patterns are shape-independent.

Finally, bacterial populations are simulated with both circuits in a scenario where cells can grow without restrictions, with an average generation time of 40 minutes. The ability of the emergent pattern to scale as the population grows was measured. Note that scaling refers to the dynamic adaptation of the pattern with size, so a scale invariant pattern would be able to maintain a constant ratio between the emergent pattern and the changing colony size^52^. Figure 10 shows representative shots of different temporal stages of the one-signal (top) and two-signal (bottom) patterns. Since the growth rate of cells is larger than the temporal scale of the signal (which is defined by the ratio between its diffusion and degradation rates), the pattern emerges during the early stages. Both patterns scale well as they are able to maintain the relative *thickness* of the green and blue regions. In the one-signal pattern, blue cells appear in the right location for the pattern to be maintained although it is blurred and discontinuous.

## DISCUSSION

Division of labor (DoL) has been defined as an evolutionary process that occurs when cooperating individuals specialize to carry out specific tasks in a distributed manner^53^. It can happen as a consequence of multicellularity but is also found in groups of similar individuals like microbes^54^. It has been theorized to require two fundamental conditions: (i) the division of tasks has to provide an inclusive fitness benefit to all of the individuals involved^10,26^ and, (ii) individuals have to slightly differ at the genetic or phenotypic level. This work has studied the conditions for the emergence and maintenance of DoL by means of the cell-based simulator *gro*.

The first hypothesis was tested and DoL was implemented through the tasks of production of two different diffusible molecules. Simulations of populations with different degrees of specialization were compared for different reward functions. Results indicate that DoL returns higher fitness benefits that individualism only if: (i) the shared molecules remained in the local proximity of the producer (short and medium communication length) and, (ii) the benefits for task specialization were accelerating. Both conclusions, although hypothesized, had never been theoretically or experimentally tested before. Therefore, these results serve as the first proof of the fitness requisites for DoL in a realistic simulated environment.

The second condition for DoL was extended, as it was proved that no genetic differences are required for cell differentiation. While it has been evidenced that phenotypic and genetic variability can lead to DoL, in this work, it was shown how DoL can also emerge as a consequence of cell communication in an isogenic group of cells. Two genetic networks that generate consensual and reversible specialization were presented and characterized. In the proposed models, cells self-organize through the exchange of a certain molecule(s) that dictates when each cell should be performing each task. Cells interact and coordinate behaviors at the local level, which ensures a precise and appropriate ratio of the different phenotypes, even in small groups. In addition, the presented models allow for the emergence of DoL without the requirements of any fitness benefits.

It was shown how DoL can be achieved transiently, where cells do not permanently specialize but still coordinate to provide global fractions of cells that perform two different tasks. This type of transient DoL could be interesting for multicellular organisms or bacterial colonies that need to adapt to sudden or temporary changes in the environment. For example, giving up the ability to reproduce momentarily in favor of helping others could be more beneficial in evolutionary terms than a permanent specialization^55^. Transient DoL also allows for the compartmentalization of different processes in a delocalized manner, which can also be more efficient in certain environments^56–58^. Depending on the processes that cells would need to complete, sensitivity thresholds and the nature of the coordinating molecule(s) could adapt to provide different proportions of individuals in each task^59^. For example, a high cost metabolic process may only be carried out by a small fraction of cells, as in the case of soma and germ cells^60^.

On the other hand, permanent DoL offers the opportunity for total specialization as a result of collective agreement. Because of this, spatial patterns of cells of different nature can emerge. This type of DoL could then be useful when tasks have to be spatially arranged or are incompatible and need to be carried out in separated locations^15^.

The two-signal circuit has proved to be more robust than the one-signal circuit. This means that the extended version is able to provide DoL for a larger parameter region. Not only that, but it also provides a permanent DoL solution for almost all conditions. The only fundamental requirement for this motif to provide a permanent solution is some noise in the cell behavior. This is easily achieved with slightly noisier production times of the regulating molecules.

To make these circuits more interesting, the permanent solutions of the one and two-signal motifs presented the formation of spatial patterns of specialized cells. The proposed regulatory mechanisms are original and lead to interesting spatial patterns that are unprecedented to this date. Both circuits are able to create *de novo* patterns that can scale with size. Both motifs present an incoherent regulation of their states, a feature also shared with other Turing-like motifs. In addition to their originality, the exceptionality of the one-signal circuit should be highlighted as it is one of the few modelled motifs able to generate patterns in growing colonies using only one regulating signal^61,62^.

The presented logic motifs were also translated into ordinary differential equations that were solved in a spatial grid. The complete set of equations and the temporal and spatial solutions can be found in the supporting information (SI2). The translation of the logical rules that an individual-based model requires to an analogous model proved to be challenging. Still, the solutions from the mathematical equations agree with the results here presented. Permanent DoL emerged as the result of signal diffusion and created similar spatial patterns for both the one and the two-signal models. This provides further proof of the robustness of the proposed mechanisms, and validates the results obtained with *gro* proving they are not a consequence of any simulation artefact.

The use of individual-based models (IbMs) like *gro* in the study of multicellular related phenomena has been greatly underused^63^. Some possible explanations include the complexity of developing robust, verified simulators, a process that is costly and time-consuming^64^. IbMs usually require a large number of parameters, where variation in their numerical values can strongly influence the resulting predictions. The precise value of some parameters should be extracted by individual measurements of the cell processes, however, IbMs are often fed with interpolated data from global phenomena. This greatly constraints the prediction capacity of the simulator. However, if one is not looking for accurate predictions of specific systems, IbMs can help providing useful information about qualitative trends as they can address evolutionary relevant questions such as the transitions from individual cells to multicellular communities^65^. A major challenge in the study of complex systems, like microbial communities, is to understand how apparently organized collective behavior emerges out of the small-scale interactions between the individuals. Complex system research requires a bottom-up approach, describing types of individuals and environments and then simulating what kind of complex dynamics emerge from the whole system. So, if one is interested in understanding processes such as the emergence of DoL or multicellularity, an IbM should be the framework of choice, since such phenomena are emergent properties that result from the collectiveness of individual interactions^66^.

## FUTURE WORK

To begin with, it would be interesting to cross-validate important results and conclusions with other similar frameworks. There are a series of related individual-cell models that were built to simulate with high detail some specific problems. For example, simulators like iDynoMiCS^67^ utilize chemical equations that allow for more realistic simulations of the diffusion of chemical fields such as nutrients or signals. Other frameworks, like CellModeller^68^ have a more precise physical engine that permits simulations of 3D environments and also cell adhesion. Replicating the experiments presented here with those frameworks could inform, for example, about the relevancy of the effect of the third dimension in bacterial groups. The cell-based Chaste software^69^ can also account for changes in cell–cell adhesion and it is ideal for dense tissue-like environments. Simulations on DoL and pattern formation utilizing this software could improve the biological relevancy of the conclusions presented here.

Secondly, the conditions favoring DoL, could be more rigorously established. Similar to the work performed by Tsoi et al.^58^, where they analyzed different metabolic pathways to derive a general criterion for outperforming DoL, the different degrees of specialization could be measured as a function of their proposed readout: the maximization of the overall productivity. It is worth mentioning that, although initial assumptions and models greatly differ, resulting conclusions offer the same principles.

Finally, implementing the genetic motifs into synthetic networks to experimentally study their potential as pattern generators could allow the development of high-level principles of biological self-organization that underlie embryogenesis in general^70–72^.

## MATERIAL AND METHODS

All examples were run with the extended version of the *gro* simulator^29^. The source code for the simulator may be found at: http://www.lia.upm.es/software/gro/. Mathematical equations (S2) were solved in Matlab by means of the ODE45 method, a versatile ODE solver that implements a variable step Runge-Kutta scheme. The spatial discretization is a 100×100 2D grid where the diffusion is modelled with a 2nd order finite difference scheme.

## ASSOCIATED CONTENT

### Supporting Information

**Text SI.** Supplementary figures

**Text SI 2.** Mathematical equations of the one and two-signal circuits and their temporal and spatial solutions.

## AUTHOR INFORMATION

### Corresponding Authors

Paula Gregorio-Godoy, Alfonso Rodríguez-Patón

### Author Contributions

The manuscript was written by P. Gregorio-Godoy. Simulations were run by P. Gregorio-Godoy and designed by G. Pérez del Pulgar and P. Gregorio-Godoy. The mathematical equations of the SI were solved by M. Rodríguez. All authors have given approval to the final version of the manuscript.

